# Beyond the myosin mesa: a potential unifying hypothesis on the underlying molecular basis of hyper-contractility caused by a majority of hypertrophic cardiomyopathy mutations

**DOI:** 10.1101/065508

**Authors:** Suman Nag, Darshan V. Trivedi, Saswata S. Sarkar, Shirley Sutton, Kathleen M. Ruppel, James A. Spudich

## Abstract

Hypertrophic cardiomyopathy (HCM), the most commonly occurring inherited cardiovascular disease, is primarily caused by mutations in human β-cardiac myosin and myosin binding protein-C. It has been thought that such mutations in myosin increase the intrinsic force of the motor, its velocity of contraction, or its ATPase activity, giving rise to hyper-contractility. We hypothesize that while these parameters are mildly affected by most myosin HCM-causing mutations, a major effect of a majority of myosin HCM mutations is likely to involve an increase in the number of myosin heads that are functionally accessible (N_a_) for interaction with actin in the sarcomere. We consider a model involving three types of interactions involving the myosin mesa and the converter domain of the myosin motor that hold myosin heads in a sequestered state, likely to be released in a graded manner as the demands on the heart increase: 1) the two myosin heads binding to one another, 2) one head binding to its own coiled-coil tail, and 3) the other head binding to myosin binding protein-C. In addition there is clear evidence of interaction between the coiled-coil tail of myosin and myosin binding protein-C. Experimentally, here we focus on myosin head binding to its own coiled-coil tail and to myosin binding protein-C. We show that phosphorylation of the myosin regulatory light chain and myosin binding protein-C weaken these respective associations, consistent with known enhancements of sarcomere function by these phosphorylations. We show that these interactions are weakened as a result of myosin HCM mutations, in a manner consistent with our structural model. Our data suggests a potential unifying hypothesis for the molecular basis of hyper-contractility caused by human hypertrophic cardiomyopathy myosin mutations, whereby the mutations give rise to an increase in the number of myosin heads that are functionally accessible for interaction with actin in the sarcomere, causing the hyper-contractility observed clinically.

## Introduction

Hypertrophic cardiomyopathy (HCM) is a heritable cardiac muscle disorder and one of the leading causes of sudden death in athletes and those under age 35 (*1, 2*). Clinically, HCM is characterized by a thickening of the ventricular heart walls and enlargement of individual cardiomyocytes (*3-5*). About half of all patients suffering from HCM are found to have mutations in genes encoding cardiac sarcomeric proteins, predominantly human β-cardiac myosin (~40%), myosin binding protein C (MyBP-C) (~40%) and the regulated thin filament system consisting of the actin-tropomyosin-troponin complex (~10%) (*3, 4, 6-9*). A leading hypothesis for the effects of myosin HCM mutations is that they increase one or more of the key parameters that determine the power output of the heart (*10*). Since power output is the product of force and velocity, the key parameters are the velocity of actin movement along an ensemble of motors, and the ensemble force produced (F_ensemble_). The latter can be estimated from F_ensemble_ = F_intrinsic_ (t_s_/t_c_) N_a_, where F_intrinsic_ is the intrinsic force of the motor, t_s_ is the time of the ATPase cycle that myosin is tightly bound to actin, t_c_ is the total cycle time of the actin-activated myosin ATPase, t_s_/t_c_ is the duty ratio, and N_a_ is the number of myosin heads that are functionally accessible for interaction with actin in the sarcomere. Measurements using mouse α-cardiac myosin as a model system showed a 2.3-fold enhancement in ATPase activity for mouse α-cardiac myosin carrying the HCM-causing R403Q mutation, a 60% increase in actin velocity, and a 40% increase in F_intrinsic_ compared to WT myosin (*11*). Similarly, R453C mouse α-cardiac myosin showed an 80% increase in F_intrinsic_ compared to WT (*11*). All of these increases would contribute to an increase in power output by the sarcomere, giving a reasonable explanation for how these mutations lead to hyper-contractility clinically. However, work by Lowey et al. (*12*) comparing the effects of the R403Q mutation in the mouse α-vs β-cardiac myosin backbone suggested a different story, and our recent studies utilizing an expression system that allows production of functional human β-cardiac myosin (*13, 14*) have confirmed that the earlier mouse a-cardiac myosin results do not accurately reflect the effects of HCM-causing mutations on the human motor.

The changes that result from either the R403Q or R453C mutations carried in the expressed human β-cardiac myosin are smaller than previously described in the mouse α-cardiac myosin model, with the largest change observed being a ~50% increase in F_intrinsic_ for R453C human β-cardiac myosin compared to wild type (*15*) (Table 1). The maximum actin-activated ATPase for R453C human β-cardiac myosin, however, was actually decreased by 30% and the velocity was decreased by 20% (*15*). In the case of R403Q human β-cardiac myosin, the maximum actin-activated ATPase was increased only 30% compared to the 2.3-fold increase seen with the mouse α-cardiac myosin (*11*), velocity was increased only 15%, and the F_intrinsic_ actually decreased by 20% (*16*) (Table 1). More recent studies of three converter domain mutations in human β-cardiac myosin, R719W, R723G and G741R, gave similar small changes in F_intrinsic_, velocity and ATPase relative to WT, with most changes being only ~10% and the largest change being ~40% (*17*) (Table 1). Thus, for all five of the HCM mutant human β-cardiac myosin proteins examined, the changes in these fundamental parameters are all less than 50%, and some parameters change in a hyper-contractile direction while others change in a hypo-contractile direction, making it difficult to determine whether there is a net hyper-contractility or a net hypocontractility caused by these HCM mutations. Thus, upon studying the human β-cardiac myosin, changes in these fundamental parameters do not appear to be able to explain the hyper-contractility that these mutations cause clinically. Something is missing.

**Table. 1.**
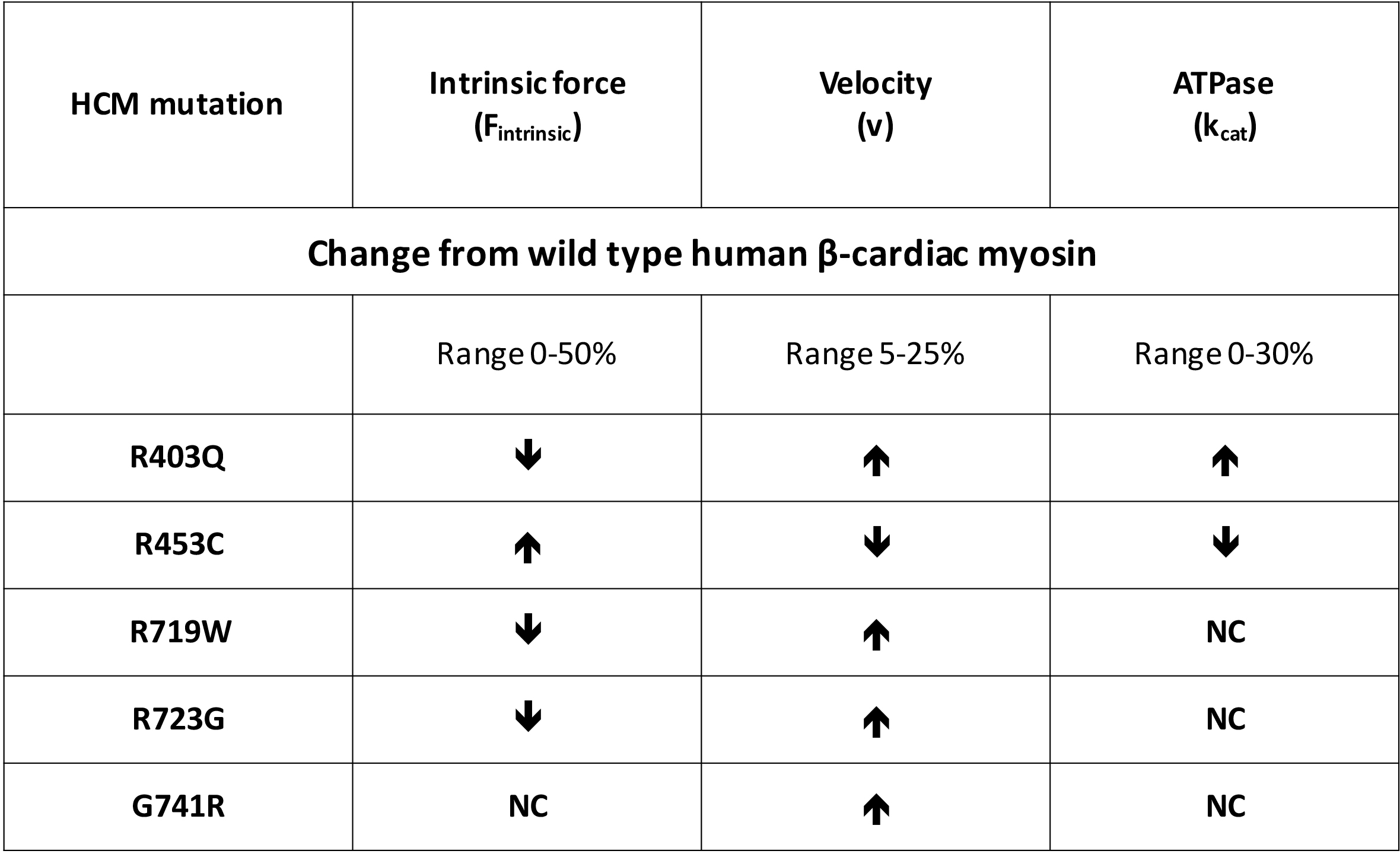
Changes in fundamental parameters of the actin-sS1 system due to 5 HCM mutations compared to WT sS1. Changes in intrinsic force, velocity and maximum ATPase for all five HCM mutant forms of human β-cardiac sS1 were found to be ≤ 50% relative to WT human β-cardiac sS1. Data for R403Q taken from Nag et al. (*16*), for R453C from Sommese et al. (*15*), and for R719W, R723G and G741R from Kawana et al. (*17*). NC = no change.

One ignored parameter in the F_ensemble_ equation is N_a_, the number of myosin heads that are functionally accessible for interaction with actin. There is growing evidence that this parameter may be key in fine-tuned regulation of the heart (*18-26*), and, as described here, may be pivotal in determining the hyper-contractility known to arise from HCM mutations in human β-cardiac myosin. Sequestered myosin heads, folded back onto the proximal part of their own α-helical coiled-coiled tail, have been reported by a number of investigators (*18, 20, 27, 28*), and Cooke and colleagues have described a super-relaxed state of myosin molecules in both skeletal and cardiac fibers that he and others have proposed to be related to this folded-back sequestered state (*22, 23*). The folded state involves an asymmetric, intramolecular interaction between the two myosin heads, first visualized by electron microscopy of dephosphorylated smooth muscle HMM (*26*). Subsequently, this folded motif was identified in electron micrographs of single molecules of myosin II isoforms from several species (*29, 30*), as well as in isolated thick filaments from tarantula skeletal (*21*), Limulus (*31*) and vertebrate cardiac muscle (*32*).

In the course of our studies, we observed that a relatively flat surface of the myosin motor domain, which we named the myosin mesa, is nearly completely conserved in all cardiac species from mouse α-cardiac myosin to human β-cardiac myosin (*33*), which is not true of other surfaces of the catalytic domain. The myosin mesa is a hotspot for myosin HCM mutations, with ~70% of the variants in the human population that map to this region being disease producing, and many of these mutations are changes from arginine residues (*33*). Only ~20% of variants in other regions of the myosin motor domain are categorized as HCM pathogenic mutations (*34*). Other hotspots are the converter domain and the proximal tail region (proximal S2; the first 126 residues of S2) of myosin (*34*).

We proposed that the cluster of arginine residues on the mesa that are mutated in HCM is likely to act as a binding domain for another protein, and binding of that protein to the mesa would hold those heads in an inactive form and thus regulate the number of myosin heads in the cardiac sarcomere that are functionally accessible (N_a_) for interaction with actin (*33*). We noted two candidate binding partners in the area of the myosin heads in the sarcomere (*33*): 1) the proximal tail region (proximal Subfragment 2, or proximal S2) of the myosin itself, and 2) myosin binding protein-C (MyBP-C), which is part of the myosin thick filaments in a ratio of 1 MyBP-C to ~6 myosin molecules (*35*). The conservation of residues on the mesa holds up when comparing fast skeletal myosin with cardiac myosin (both muscle types have MyBP-C), while the conservation is lost when comparing smooth muscle myosin or non-muscle myosin II with cardiac myosin (neither smooth muscle or non-muscle cells have MyBP-C) (*33*). This supported our idea that MyBP-C might be a possible binding partner.

Here we directly demonstrate the binding to S2 of both short Subfragment 1 (sS1), a truncated myosin head containing the essential light chain (ELC) but not the regulatory light chain (RLC), and of a double-headed myosin construct containing both the ELC and the RLC. We extend our studies to show that the RLC phosphorylation weakens this interaction. Additionally, we also show that the sS1 myosin binds to MyBP-C and this interaction is weakened by the phosphorylation of the M domain. Finally, we discuss a potential unifying hypothesis for the molecular basis of hyper-contractility caused by a majority of human hypertrophic cardiomyopathy myosin mutations, whereby the mutations give rise to an increase in the number of myosin heads that are functionally accessible for interaction with actin in the sarcomere, causing the hyper-contractility observed clinically.

## Results

### A model of a sequestered complex of human β-cardiac myosin highlights potential electrostatic interactions between HCM-involved arginine residues on the myosin mesa and HCM-involved aspartate/glutamic residues on the proximal part of S2

It was previously proposed that the myosin mesa on the globular S1 head of human β-cardiac myosin might interact with its own S2 coiled-coil tail and/or with myosin binding protein-C (MyBP-C), sequestering the myosin heads in a state that is not accessible for interaction with actin (*33*). Indeed, a number of different myosin isoforms have been shown to adopt a compact structure where the two S1 heads are folded back onto their own proximal S2 tail (Fig. 1A)(*18-21, 26*). Fig. 1 shows a homology-modeled folded-back structure of human β-cardiac myosin from the 3D-reconstructed images of tarantula skeletal muscle thick filaments by Alamo et al. (*19*). One head (Fig. 1B, left) is called the ‘blocked head’ (outlined with a black line) since its actin-binding domain is not available for interaction with actin, and the second head is called the ‘free head’ (Fig. 1B, right) since its actin binding domain is still available (*26*). Fig. 1C shows the blocked head (with its ELC colored brown, and the RLC green) on the left interacting with the proximal S2.

**Figure 1.**
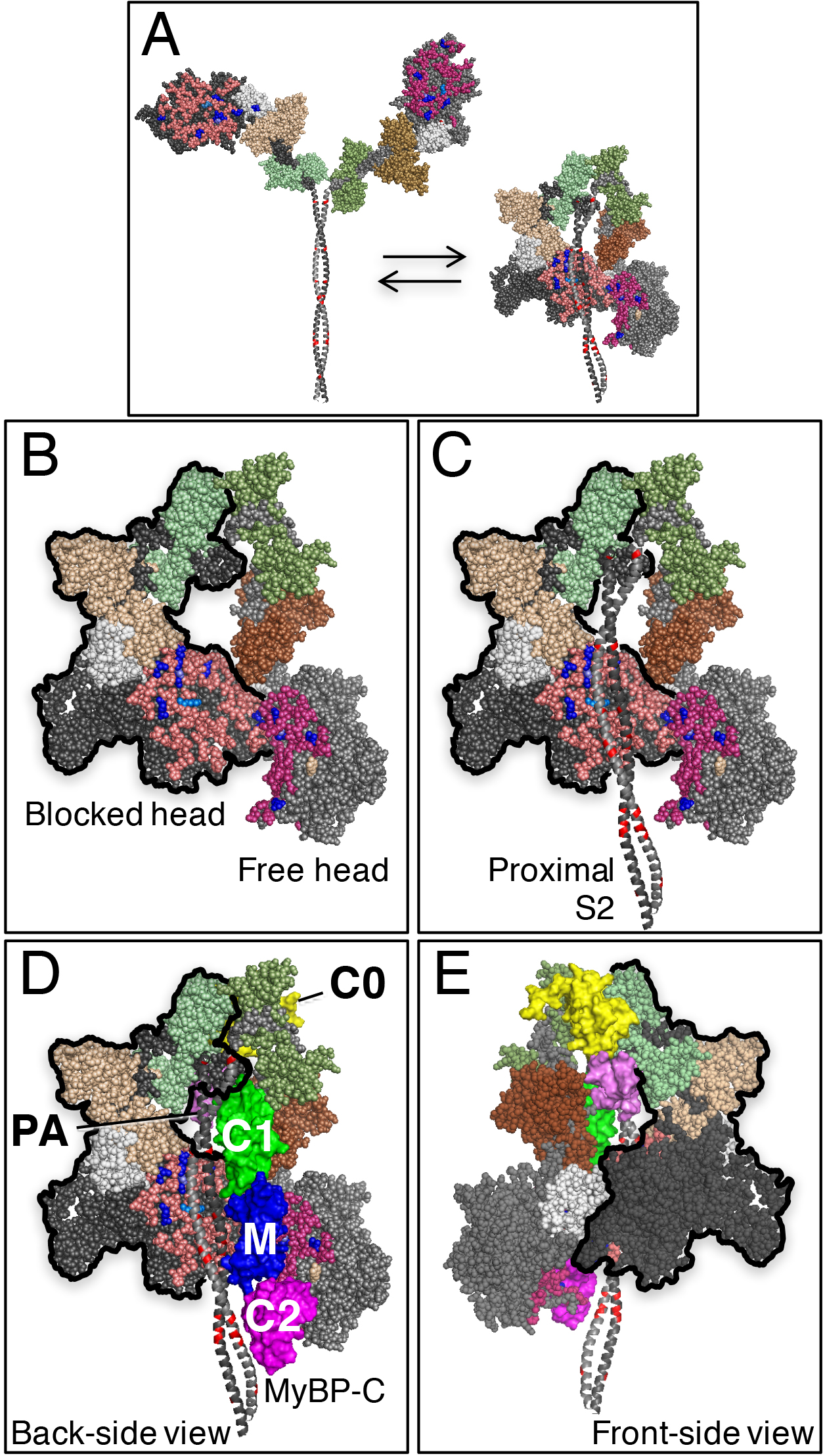
Structural model of sequestered heads of human β-cardiac myosin based on the 3D-reconstruction of tarantula skeletal myosin thick filaments by Alamo et al. ( *19*) (PDB 3DTP). **A.** A short version of myosin HMM, showing only 126 residues of the coiled-coil S2 domain, is illustrated in its open and closed states, which are in equilibrium. The heavy chain residues of the S1 head on the left are colored pink (mesa residues), dark blue (arginine HCM mutations), light blue (a lysine HCM mutation), white (the converter domain), and dark grey (all remaining residues). The ELC is colored light brown and the RLC is light green. The heavy chain residues of the S1 head on the right are colored dark pink (mesa residues), dark blue (arginine HCM mutations), light blue (a lysine HCM mutation), white (the converter domain), and light grey grey (all remaining residues). The ELC is colored dark brown and the RLC is dark green. The positions of glutamate and aspartate residue HCM mutations in the proximal S2 tail are shown in red. **B.** The back-side view (named from the 3D-reconstruction of the tarantula thick filament; this side faces the myosin bipolar thick filament) of the sequestered state showing the potential interaction between different domains of the two heads. **C.** The back-side view of the sequestered state showing the myosin mesa domains of the two heads cradling the proximal S2, with potential interactions between the proximal S2 and the mesa of the blocked head (on the left, outlined with a black line). **D.** The back-side view of the sequestered state showing the myosin mesa domains of the two heads cradling the C0-C2 domains of MyBP-C, with potential interactions between the C0-C2 and the mesa of the free head (on the right) and illustrating potential interactions between proximal S2 and C1-C2 (*37*). The yellow C0 domain is bound to the RLCs (*36*) on the front-side view of the complex and the PA domain (pink) connects to the C1 (green)-M (blue)-C2 (magenta) domains, which are on this back-side view of the complex. **E.** The front-side view of the complete complex showing the blocked head (on the right, outlined with a black line) interacting with the converter domain (white) of the free head (on the left).

Furthermore, in the folded structure we have positioned the N-terminal C0 domain (yellow) of myosin binding protein-C (MyBP-C) bound to the two RLCs (Fig. 1D), which has been shown to occur experimentally (*36*). Similarly, we have positioned the C1-C2 domains of MyBP-C in the folded structure bound to the first 126 residues of the myosin S2 tail (proximal S2) (Fig. 1D), which also has been shown to occur experimentally (*37*).

When the structure is viewed from its back side (the side facing the shaft of the thick filament in the 3D reconstructions (*19*) (Fig. 1C), several observations can be made. Strikingly, the proximal S2 is cradled by the mesa domains of the S1 heads (light pink residues on the blocked head and dark pink residues on the free head). Also striking is that a cluster of 6 HCM arginine mutations on the blocked head mesa (Fig. 1C, blue residues, R169G, R249Q, R403Q, R442C, R453C and R663H) and 1 HCM lysine mutation (light blue residue, K657Q) are in close proximity to a cluster of 8 HCM glutamate/aspartate residues on the proximal S2 (Fig. 1C, red residues, E875del, E894G, E903G, D906G, E924K, D928N, E930K, E931del), suggesting there may be an electrostatic interaction that is important for functioning of the sarcomere and therefore sensitive to changes in any of these residues. Note that the S2 tail in this model is in closest contact with the mesa of the blocked head (Fig. 1C). This leaves space for the potential binding of the MyBP-C C1-C2 region to both S2 and to the mesa of the free head (Fig. 1D).

When the structure in Fig. 1D is rotated 180° about its vertical axis to reveal the front side, the surface adjacent to the actin-binding surface and the mesa of the blocked head can be seen to bind to the converter domain (Fig. 1E, white residues) of the free head. It is interesting to note that there are virtually no hypertrophic cardiomyopathy (HCM) mutations on the surfaces of the S1 catalytic domains seen from this front view (*34*)

Based on these observations, we propose the existence of a sequestered complex of human β-cardiac myosin S1 heads in which the heads are not available for interaction with actin, and which may be held together by four types of interactions, each of which individually may be relatively weak: S1-S1 (Fig. 1B), S1-S2 (Fig. 1C), S1-MyBP-C (Fig. 1D), and S2-MyBP-C (Fig. 1D).

Here we primarily focus on a here-to-fore unknown interaction of the myosin head with S2, which is weakened by RLC phosphorylation. Furthermore, we show that the effects of various HCM mutations on the affinity of the S1-S2 interactions are in keeping with the model shown in Fig. 1. We also present evidence of the binding of both full length MyBP-C and the N-terminal fragment C0-C2 to sS1, binding that is weakened by phosphorylation of the M-domain in each case. Finally, we present arguments suggesting that many, if not most, myosin HCM mutations are effectively weakening this sequestered complex, liberating more heads into play during systolic contraction, which results in hyper-contractility of the heart.

### Human β-cardiac short S1 binds to the proximal part of human cardiac S2

The folded-back structures found by 3D-reconstructions of tarantula skeletal and human cardiac myosin thick filaments and other myosin structures (*18-21, 27, 28, 38-42*) are generally described as being stabilized by an interaction between the two myosin heads (Fig. 1), but the above considerations encouraged us to look for interactions between sS1 and the proximal S2. Since S1 and S2 are connected in the intact myosin molecule, we estimated the effective concentrations of S1 and S2 in the intact molecule to be ~60 µM (see Supplemental materials). Thus, at 100 mM KCl (approaching physiological ionic strength), a K_d_ even in the double-digit µM range may be physiologically relevant. Microscale thermophoresis (MST), which follows the diffusion of a fluorescent probe along a temperature gradient, can be used to measure bio-molecular interactions over a broad range of affinities, including a K_d_ in the double-digit µM range (*43-45*).

After demonstrating that the MST approach validly measures the affinities of four known protein-protein interactions involving sS1, actin, proximal S2 and MyBP-C (Fig. 2A), we used sS1 fluorescently tagged with Cy5-labeled cysteine residues (Fig. 2B, open circles) or with a C-terminal GFP (Fig. 2B, closed circles) to measure binding to proximal S2. In both cases we observed binding between sS1 and proximal S2 with a K _d_ of 3-5 µM in buffer containing 25 mM KCl (Fig. 2B orange curves). This affinity became weaker (30-70 µM, Fig. 2B black curves) by increasing the salt concentration to 100 mM KCl, indicating an electrostatic interaction between the sS1 and proximal S2. A control with GFP alone showed no binding to S2 (Fig. 2B, green curve).

**Figure 2.**
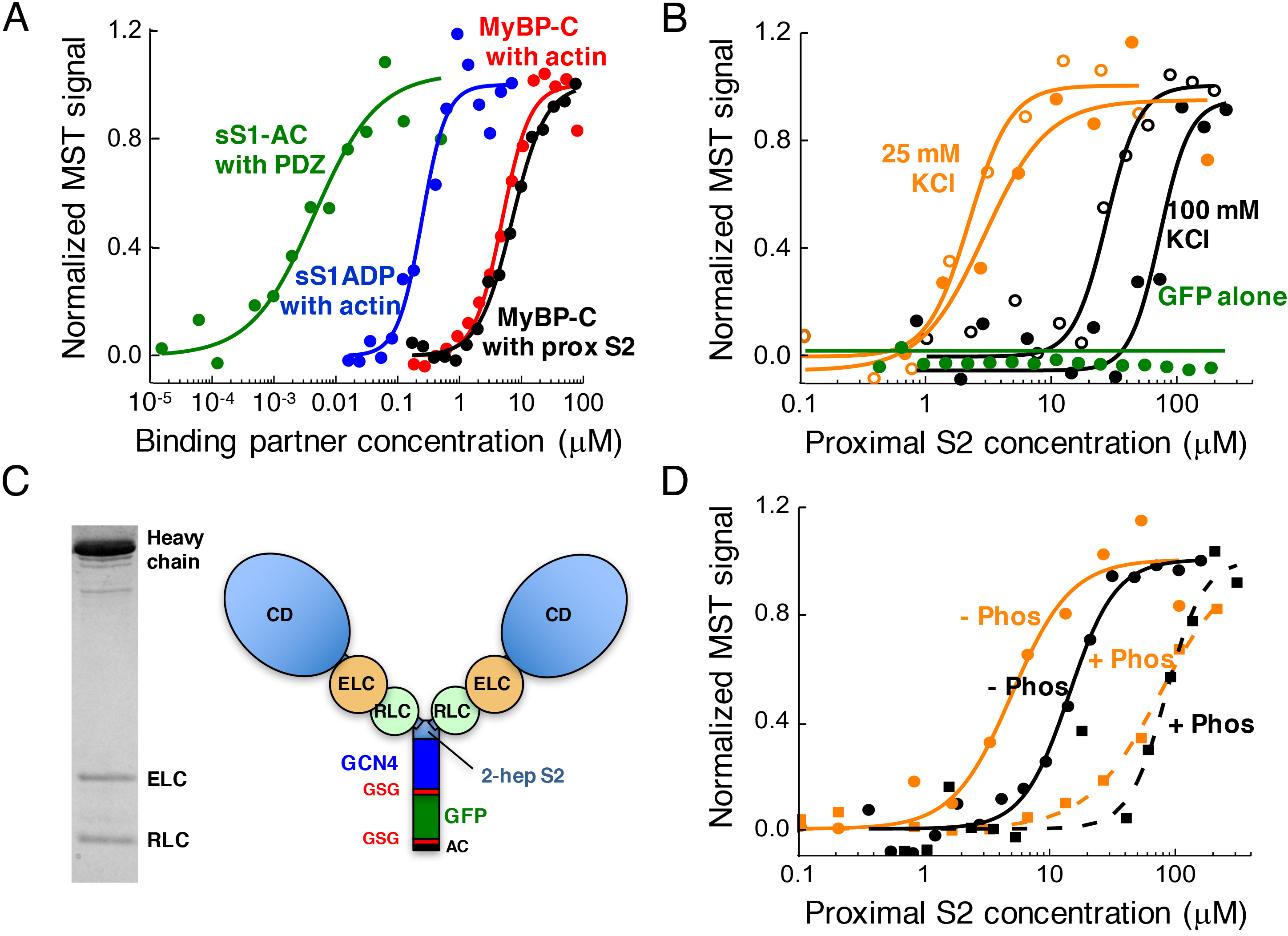
Binding of sS1 and 2-hep HMM to proximal S2 using MST. **A**. MST technology gives the expected K_d_ for four protein-protein interactions known from earlier studies using other techniques: sS1 with a C-terminal 8-residue affinity-tag (RGSIDTWV) (sS1-AC), which binds to PDZ (used for binding of sS1-AC to a PDZ domain for in vitro motility measurements) (~ 5 nM, green) (*59*); sS1 binding to actin in the ADP state (~250 nM, blue) (*16, 60*); MyBP-C binding to actin (~5 uM, red) (*61*), and MyBP-C binding to proximal S2 (~7 uM, black) (*37*). In all cases, experiments were performed at 23^°^C and the measured values are within 2-fold from published values. **B**. Binding of Cy5-labeled (open circles) and GFP-tagged (closed circles) sS1 to proximal S2 at 25 mM (orange) and 100 mM (black) KCl. Throughout the manuscript GFP refers to the enhanced version (eGFP) of GFP. Increasing concentrations of proximal S2 were added to 50 nM WT sS1, and the fluorescently-labeled sS1 was followed to observe changes in diffusion rate. GFP alone showed no binding (green curve). **C**. SDS PAGE of 2-hep HMM and a schematic drawing of its structure. The two S1 heads are linked by 2 heptad repeats of the S2 region followed by a GCN4 leucine zipper to ensure dimerization. Following the GCN4 is a GSG swivel linker, a GFP, another GSG linker, and ending in the 8-residue (RGSIDTWV) affinity clamp (AC). **D**. Binding of GFP-labeled 2-hep HMM to proximal S2 at 25 mM (orange) and 100 mM (black) KCl. De-phosphorylated (-Phos) and phosphorylated (+Phos) 2-hep HMM are compared.

### Dephosphorylated human β-cardiac short HMM binds to the proximal part of S2, and phosphorylation of the myosin RLC weakens the affinity of the binding

Phosphorylation of the human β-cardiac RLC serine15 residue has significant effects on cardiac muscle force production, shortening velocity, and peak power output, and is altered during cardiac disease states (*46, 47*). In order to explore the effects of RLC phosphorylation on the binding of the S1 head to S2, we created a 2-headed human β-cardiac myosin construct that resembles the conventional fragment of myosin called heavy meromyosin (HMM). Like HMM, this construct is a soluble monomeric species that consist of 2 heads connected by a coiled-coil tail. It consists of the human β-cardiac myosin heavy chain (HC) residues 1-855, containing both the human cardiac ELC and RLC binding sites and only 14 residues (2 heptad repeats) of the N-terminal part of S2 (Fig. 2C). A GCN4 leucine zipper was inserted to ensure dimerization (*48*) and GFP was placed near the C-terminus to follow its fluorescence in the MST binding experiments. This construct, referred to as 2-hep HMM throughout, shows excellent activity in the in vitro motility assay, propelling actin filaments at 1.5 µm s^-^1 at 23^°^C, and an intrinsic force using the laser trap single molecule assay of 1.8 pN. The stoichiometry of the complex was 1:1:1 for HC:ELC:RLC (see Materials in Supplementary materials).

Evidence supports the idea that RLC phosphorylation liberates more myosin heads into an active form in cardiac muscle (*24*). We therefore used the MST binding assay to test whether the binding of the human β-cardiac 2-hep HMM to S2 is weakened by phosphorylation of its RLC (Fig. 2D). The RLC on the purified 2-hep HMM was fully de-phosphorylated. To obtain the phosphorylated form, we treated the 2-hep HMM with Ca^2+^-CaM-activated MLCK and ATP, and confirmed that more than 90% of the RLC was phosphorylated by urea-SDS-glycerol gel electrophoresis (*49*). While the dephosphorylated 2-hep HMM bound proximal S2 with a K_d_ of 5 µM at 25 mM KCl (Fig. 2D, solid orange curve), the phosphorylated 2-hep HMM bound with a much lower affinity (K_d_ = 70 µM, Fig. 2D, dashed orange curve). At 100 mM KCl, the K_d_ for de-phosphorylated 2-hep HMM was 15 µM (Fig. 2D, solid black curve), while the phosphorylated 2-hep HMM again bound with a much lower affinity (K_d_ = 90 µM Fig. 2D, dashed black curve).

### The effects of human β-cardiac myosin HCM mutations on the affinity of the sS1-S2 binding are consistent with the hypothesized sequestered complex

If the myosin mesa interacts with proximal S2, thereby sequestering myosin heads, we hypothesize that the S1 mesa arginine/lysine residues and the proximal S2 glutamate/aspartate residues that give rise to HCM, when mutated, shift the equilibrium of the S1-S2 interaction to a more open state. This would increase the number of functionally-accessible heads, causing hyper-contractility of the sarcomere, consistent with clinical observations. To this end we investigated the binding of the HCM-causing R453C human β-cardiac sS1 to proximal S2, since this mutation lies very near to proximal S2 in the sequestered complex model (Fig. 3A). In all studies reported here we compared the mutant proteins to a WT control at the same time to control for variabilities in conditions from day to day. Both sS1-GFP and sS1-Cy5 carrying the R453C mutation showed weaker binding affinities for proximal S2 than their WT sS1 counterparts. Fig. 3B shows sS1-GFP binding to proximal S2. The R453C human β-cardiac myosin (blue curve) binds much more weakly than the WT protein, suggesting that the myosin mesa is potentially involved in binding to proximal S2 and that the effect of this mesa HCM mutation may be to weaken the sequestered complex, which would allow more myosin heads to enter the pool of actin-interacting force-producing heads, and thereby increasing the contractility of the muscle. R403Q, which is on the far outer edge of the blocked head mesa, and buried in the interaction between the two heads (Fig. 3A), would not be predicted to affect the sS1-S2 interaction according to the structural model. The R403Q human β-cardiac sS1 (Fig. 3B, red curve) showed the same affinity for the sS1-S2 interaction as WT human β-cardiac sS1 (Fig. 3B, black curve). We hypothesize that the R403Q mutation is weakening one of the other three types of interactions that hold the sequestered complex together. Thus, in the model shown in Fig. 1, R403Q is in a position to possibly affect the S1-S1 interactions and/or the S1-MyBP-C interactions.

**Figure 3.**
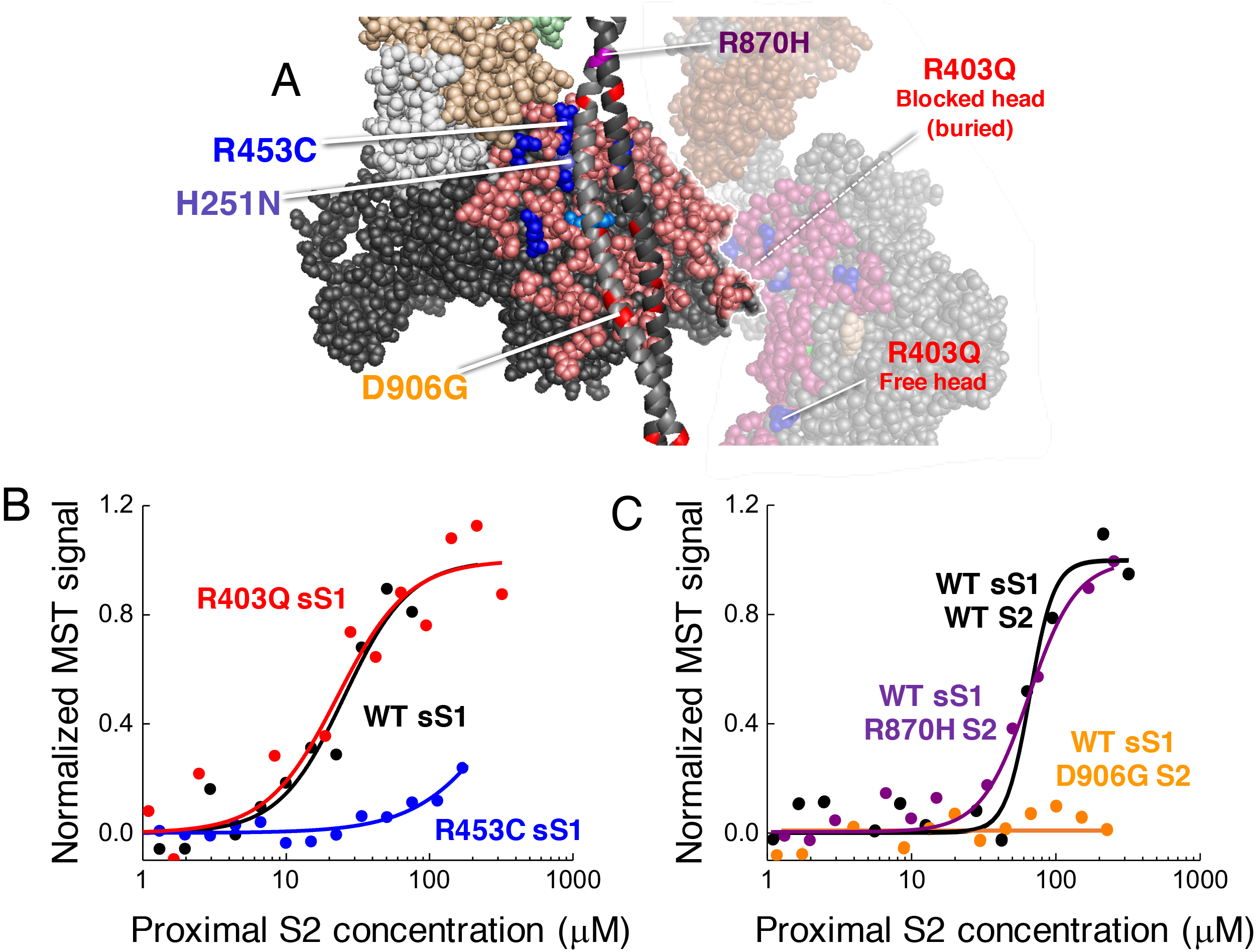
Effects of HCM mutations on the interaction of human β-cardiac sS1 with proximal S2. Binding was measured using MST in 100 mM KCl. **A**. Model of sS1 of the blocked head interacting with proximal S2, marking the positions of two sS1 HCM mutations and two proximal S2 mutations that were used for binding experiments. Also shown is the free head (translucent head) interacting with the blocked head. R453 of sS1 lies on the mesa domain and is predicted to interact with the proximal S2, whereas R403 is deep buried and away from the interacting site. Similarly, D906 on the proximal S2 is predicted to be interacting with the mesa, but R870 is not. **B**. Binding of GFP-labeled WT (black), R403Q (red) and R453C (blue) sS1 to proximal S2. **C**. Binding of GFP-labeled WT sS1 to WT proximal S2 (black), and R870H (purple) and D906G (orange) mutant proximal S2.

In a parallel study involving two early onset HCM human β-cardiac myosin mutations, Adhikari et al. (*50*) found that the H251N mutation, which is next to R453C on the mesa (Fig. 3A) in potential contact with proximal S2 in the sequestered model, also significantly weakens the binding of sS1 to S2, while the D239N mutation, which is on the other side of the molecule from the mesa, has no effect on the affinity of sS1 for proximal S2 compared to WT sS1, further bolstering the overall hypothesis presented here.

To further test our hypothesis, we examined the aspartate/glutamate HCM mutations in the proximal S2 domain that are near the mesa to see whether they also weaken the binding of sS1 to S2. Fig. 3C shows that D906G S2 (orange curve), which overlays the blocked head mesa, shows no binding to sS1 up to 250 µM proximal S2, in contrast to WT S2 (black curve). A control for the lack of binding of the D906G S2 is the proximal S2 HCM mutation R870H, which is outside the range of the mesa (Fig. 3A). R870H S2 and WT S2 binding to sS1 show the same affinity (Fig. 3C). In keeping with our hypothesis that a majority of HCM mutations weaken the affinity of one of the four types of intra- and intermolecular interactions that hold the folded sequestered complex together, the R870H proximal S2 HCM mutation has been shown by Gruen and Gautel (*37*) to decrease the affinity of binding of the proximal S2 for the C1-C2 domain of MyBP-C by more than an order of magnitude. This result could be expected by the structural model of Fig. 1, which shows R870 potentially interacting with C1 of MyBP-C (compare Fig. 1D with 3A).

### WT human β-cardiac sS1 binds to both full-length human cardiac myosin binding protein-C and to the N–terminal C0-C2 fragment, and this affinity is weakened by protein kinase A phosphorylation of the M-domain

Given the work of Gautel and colleagues showing binding of C1-C2 of MyBP-C to proximal S2 (*37*) and the model shown in Fig. 1, we hypothesized that the myosin mesa may serve as an interacting platform for MyBP-C (*33*). Both the globular head domain of β-cardiac myosin and MyBP-C are hot spots for HCM pathogenic mutations, lending support to the hypothesis that these two proteins may be interacting with one another.

Human cardiac MyBP-C, homology modeled from known structures of some of the domains (see Methods in Supplementary materials), is shown in Fig. 4A. When expressed and purified from Sf9 cells (see Methods in Supplementary materials), the protein is fully de-phosphorylated (Fig. 4A). We treated the MyBP-C with protein kinase A (PKA) and ATP, which phosphorylates the MyBP-C on the M domain (Fig. 4A, D, blue domain adjacent to the mesa of the free head).

**Figure 4.**
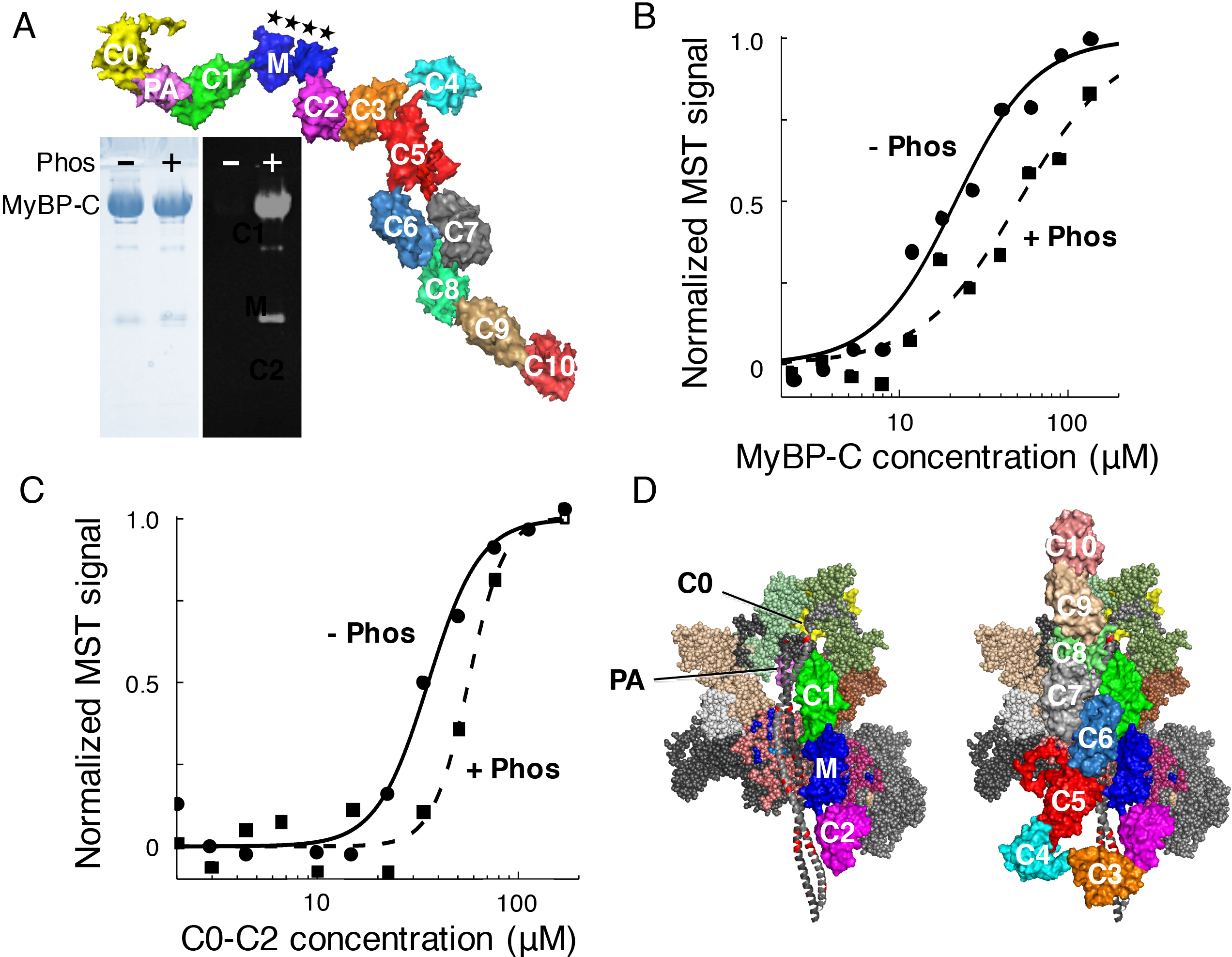
Binding of human β-cardiac sS1 to full-length human cardiac MyBP-C. **A**. Surface rendition of full-length human cardiac MyBP-C homology-modeled from known structures of C0, C1, C2, C3, C5 and M2 domains, which were obtained individually using structural homologues from their respective PDB files (see Methods). The other domain structures (C4, C6, C7, C8, C9, C10, M1, PA loop) were modeled independently using ab-initio and template-based prediction methods (see Methods). The C0-C10 domains were connected using PyMol C-term to N-term to obtain the image shown. C4 is the only domain that has its N-term and C-term emanating from the same side of the molecule, which induces a sharp bend in the molecule at the C4 position. There are four serine phosphorylation sites (★) on the M domain (blue) that regulate MyBP-C function (*62*). The left panel of the gel shows a Coomassie gel of full-length human cardiac MyBP-C purified from Sf9 cells (see Methods) treated with Lambda phosphatase (-) or with PKA and ATP (+). The right panel shows ProQ Diamond staining of the same gel. Lower molecular weight contaminants are also phosphorylated (probably proteolytic fragments of MyBP-C). **B**. Binding of Cy5-labeled sS1 to de-phosphorylated (-Phos; solid circles with solid curve) and phosphorylated (+Phos; solid squares with dashed curve) full-length MyBP-C. **C**. Binding of Cy5-labeled sS1 to de-phosphorylated (-Phos; solid circles with solid curve) and phosphorylated (+Phos; solid squares with dashed curve) C0-C2. D. Hypothetical models of the interaction of C0-C2 and full-length MyBP-C to folded-back, sequestered S1 heads. These structures are working models for experiments going forward. All domains of MyBP-C are marked. Binding experiments were measured using MST in 100 mM KCl at 23^°^C.

We tested whether de-phosphorylated MyBP-C binds to sS1, which does not contain the RLC that Ratti et al. showed binds to C0 of MyBP-C (*36*). Using MST, we found that full-length human de-phosphorylated MyBP-C binds to sS1 with a K_d_ of ~20 µM (Fig. 4B, solid curve). Phosphorylation of the MyBP-C with PKA and ATP reduced the affinity of the binding significantly (K_d_ ~50 µM, Fig. 4B dashed curve).

Very similar binding (K_d_ = ~35 µM) was seen with the de-phosphorylated C0-C2 domain alone (Fig. 4C, solid curve), consistent with the model of the complex shown in Fig. 1D and 4D. This binding was weaker after phosphorylation of the C0-C2 with PKA and ATP (Fig. 4C, dashed curve). The slightly weaker binding of C0-C2 compared to full-length MyBP-C has been consistent in duplicate experiments from multiple protein preparations and might suggest that the C4-C10 domains play some role in the sequestered complex formation. Interestingly, if one continues to connect each of the C3 to C10 domains from N- to C-terminus, the complex makes a sharp bend at C4 due to the emergence of the N- and C-termini of C4 from the same side of the C4 domain. This could result in the folding back of C3-C7 onto the blocked head mesa with C8-C10 free to interact with the thick filament backbone, as illustrated in Fig. 4D. While this is a highly speculative structure, it would explain a somewhat stronger binding of full length MyBPC to sS1 compared to the C0-C2 fragment alone, and is consistent with all known literature on the sequestered state and a possible role of MyBP-C in myosin interactions.

### Modeling of the folded-back structure shows that nearly all converter HCM mutations lie at the interface of the two folded-back S1 heads

Gautel and colleagues demonstrated binding of C1-C2 of MyBP-C to proximal S2 (*37*), and here we demonstrate the binding of sS1 to both proximal S2, MyBP-C and its C0-C2 fragment, along with the binding of 2-hep HMM to proximal S2. Besides these three types of interactions, there is clearly an S1-S1 interaction, which has been one focus of discussions by those who described folded-back structures of myosin by electron microscopy (*20, 28*). This S1-S1 interaction involves the association of a surface adjacent to the mesa of the blocked head (Fig. 5A,B, dark grey residues) with the converter domain of the free head (Fig. 5A,B, white residues). The converter domain is of great interest in HCM because it is the hottest spot for HCM mutations, where all reported variants to date in the human population prove to be HCM-causing pathogenic mutations (*34*). We therefore modeled in the converter HCM mutations into the homology-modeled folded-back human β-cardiac myosin shown in Figure 1. When examining the positions of these HCM mutations, one obtains a striking result. Nine out of ten of those mutations (only L749Q is buried) are at the interface of the free head converter and the blocked head binding face (Fig. 5). Furthermore, the HCM residue R719W on the converter (Fig. 5D,E, middle cyan residue) of the free head is in direct contact with the severe early onset HCM mutant residue D382Y (*51, 52*) on the binding face of the blocked head (Fig. 5E, red residue in spheres), possibly forming a salt bridge.

**Figure 5.**
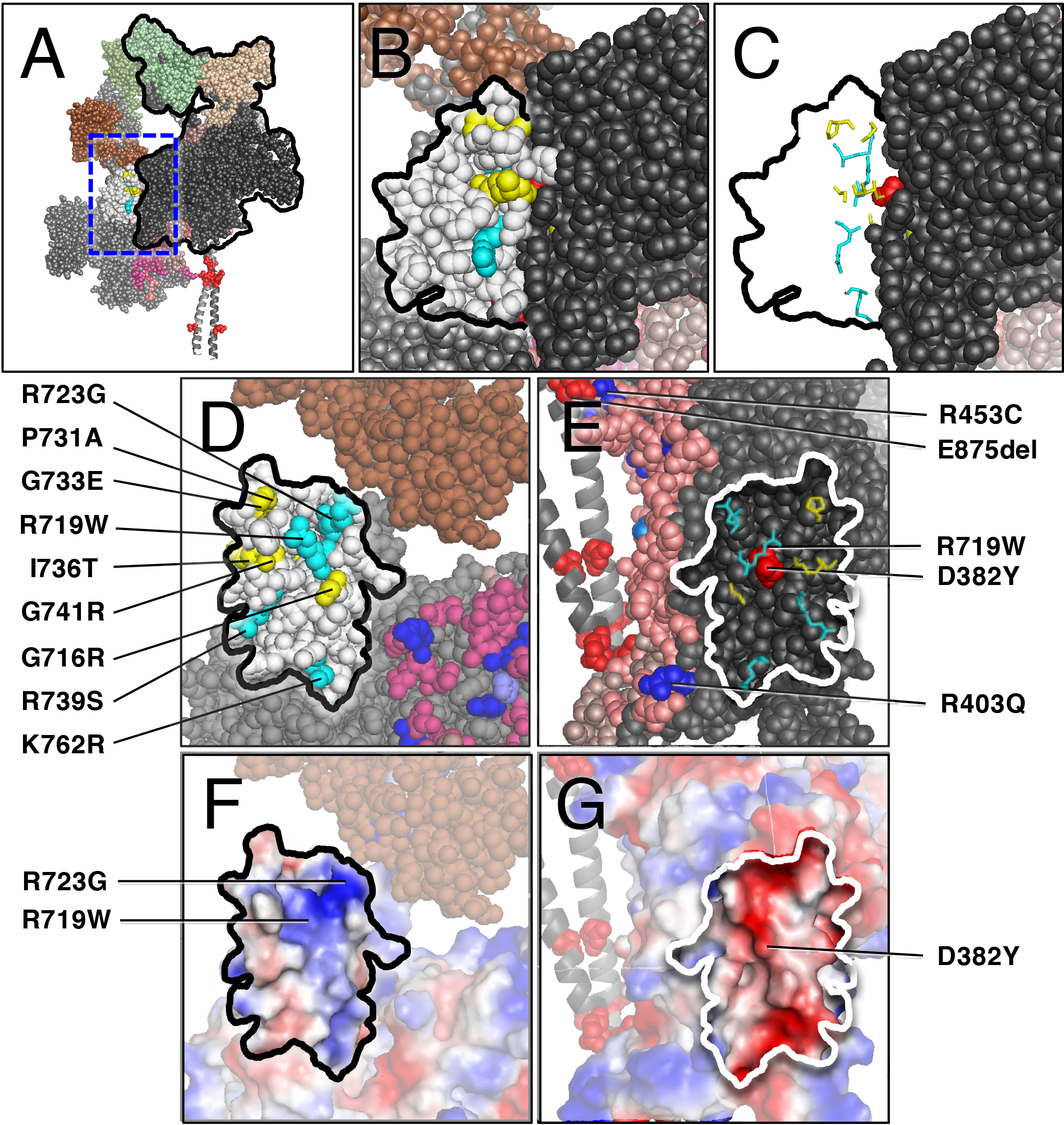
Structural features of the S1-S1 interaction in the homology-modeled human β-cardiac S1 sequestered model suggest a role of most converter mutations in weakening the affinity of the sequestered complex. **A**. The front view of the sequestered complex shows the association of a surface adjacent to the mesa of the blocked head (black outline) with the converter domain of the free head (white residues). **B**. The area outlined by the dashed blue box in A is blown up to focus on the interface between the blocked head (dark grey residues) and the converter (white residues, black outline) of the free head. Converter mutations are colored light blue for arginine and lysine residues, and yellow for uncharged residues. **C**. The 9 HCM residues shown in stick format with the remainder of the converter domain residues removed. Note the striking alignment along the binding face of the 2 S1 heads. **D**. The image in C rotated 90° clockwise about the vertical axis defining the binding interface. All 9 HCM residues are on the surface of the converter at the S1-S1 junction. **E**. The image in C rotated 90° counterclockwise about the vertical axis defining the binding interface. The positions of the 9 HCM residues of the converter domain of the free head are mapped onto the black binding interface of the blocked head. Note that R719W of the converter is in direct contact with D382Y of the blocked head surface. D382Y is an HCM mutation that causes early onset disease (*51*). R403Q is at the corner of the blocked head binding face and could be involved in the S1-S1 interaction. **F**. The same image as D, but shown in vacuum electrostatics mode in Pymol. The converter binding interface is generally positively charged. **G**. The same image as E, but shown in vacuum electrostatics mode in Pymol. The blocked head binding interface is generally negatively charged.

Two additional points can be made when viewing Fig. 5E. First, this view of R453 on the mesa and E875 on S2 show how close in proximity they are to one another, possibly forming a salt bridge. This view is a ~90° rotation to the left of the image shown in Fig. 1D, with the mesa residues colored pink in both cases. Also of note is the position of the R403Q HCM mutation (Fig. 5E), which appears to be part of the S1-S1 binding interface. Examination of the binding interface by the vacuum electrostatics mode in Pymol suggests an electrostatic interaction, with the binding interface of the converter domain of the free head being generally positively charged (Fig. 5F), with R723 and R719 being key residues, and the binding interface of the blocked head domain being generally negatively charged (Fig. 5G).

## Discussion

Before embarking on our binding studies, we verified the use of MST technology by showing that the technique gives the expected K_d_ for four known protein-protein interactions. MST has many advantages over other binding measurement techniques. Unlike antibody pull-down and sedimentation approaches, in MST one is measuring a true equilibrium not perturbed by separation of the protein complex from the unbound proteins. Furthermore, the experiments are rapid and only small quantities of proteins are needed.

The decrease in K_d_ for sS1 binding to proximal S2 when comparing R453C human β-cardiac sS1 to WT is an order of magnitude, which on its own would be predicted to release a substantial fraction of heads in the sequestered complex for interaction with actin (see Supplemental materials). If a significant portion of the myosin heads were in the sequestered state under normal conditions of heart function, such a large change may not be tolerated. Our hypothesis, however, is that in addition to the sS1-proximal S2 interaction, the sequestered complex is held together by three other associations: S1-S1 interactions, which we predict are weakened by the majority of converter HCM mutations and possibly the R403Q mutation (Fig. 5E); S1-MyBP-C interactions, which we predict are weakened by MyBP-C HCM mutations, possibly the R403Q mutation in the free head (compare Fig. 1D and 3A) and other myosin mesa HCM mutations on either head; and proximal S2-MyBP-C interactions, which we predict are weakened by proximal S2 HCM mutations (as already demonstrated for R870H and E924K by Gruen and Gautel (*37*) and by MyBP-C HCM mutations. Thus, an order of magnitude weakening or greater of any one of these, as in the cases for the mesa HCM mutations R453C and D906G reported here, the mesa mutation H251N (*50*), and the proximal S2 HCM mutations R870H and E924K reported by Gruen and Gautel (*37*), would lead to a smaller change in the fraction of heads released than expected if only one type of intra- or inter-molecular interaction were occurring.

The data presented here provide experimental support for the myosin mesa hypothesis (*33*), which submits that the highly-conserved mesa is a hotspot for HCM mutations that are predominantly weakening the stability of a sequestered complex involving interactions of the mesa with either the myosin S2 tail and/or MyBP-C, resulting in more myosin heads becoming available for interaction with actin, causing the hyper-contractility observed clinically. Furthermore, we provide evidence that phosphorylation of the myosin RLC with MLCK and of the M-domain of MyBP-C by PKA weaken the putative sequestered complex.

The idea of sequestered myosin heads, released upon phosphorylation of the myosin RLC and of the M-domain of MyBP-C, is not a new one. As early as 1999, Gruen and Gautel (*37*) showed that the C1-C2 domain of MyBP-C binds to proximal S2 and this binding is weakened and eliminated by the S2 HCM mutations R870H and E924K, respectively. In 2001, Wendt et al. (*26*) demonstrated that dephosphorylated smooth muscle HMM has its heads folded back onto its S2 tail with an asymmetric S1-S1 interaction and defined the ‘blocked’ and ‘free’ states in this folded state, and Levine et al. showed that PKA phosphorylation ‘loosens’ the myosin heads from their more organized (sequestered?) state on thick filaments (*53*). The folded HMM structure of Wendt et al. (*26*) was then described in a number of 3D reconstructions from electron micrographs of isolated muscle thick filaments in their relaxed state (*21, 31, 32*). These folded myosin structures are proposed to be the structural basis for the super-relaxed state of skeletal and cardiac myosin described by Cooke and colleagues (*22, 23*), which release nucleotide from the myosin active site extremely slowly. More recently, these ideas have received additional support, with reports that unphosphorylated cardiac MyBP-C regulates the binding of myosin to actin and significantly decelerates the ATP-induced dissociation of myosin from actin (*54*), that MyBP-C inhibits myosin thick filament activity in heart muscle by holding heads in a state parallel to the thick filament axis (*42*), that cardiac MyBP-C and its phosphorylation state regulate the rate and force of contraction in mammalian myocardium (*55*), and that RLC phosphorylation controls myosin head conformation in cardiac muscle (*24*). Furthermore, Linari et al. (*56*) recently demonstrated that at higher loads, skeletal muscle thick filaments undergo a stress-dependent transition that opens up additional myosin heads bound to the thick filament, which are recruited to aid the high-load contraction. Such a mechanism might also operate in the heart, and represents an additional level of natural regulation of cardiac contractility.

What is new here is the realization that the three hotspots for myosin HCM mutations (*34*), the myosin mesa, proximal S2, and the converter domain, are all integral parts of the folded-back sequestered complex defined by 3D-reconstructions of muscle myosins, and we hypothesize that a majority of myosin HCM mutations are likely to be weakening this sequestered state, resulting in more functionally accessible heads for actin interaction, and causing the hyper-contractility observed for HCM clinically.

Thus, we propose that the sequestered state is held together by a series of relatively weak interactions between the two S1 heads, S1 and proximal S2, MyBP-C and S1, and MyBP-C and S2. In addition, there also must be interactions between the folded heads and the light meromyosin (LMM) backbone, since the folded complex is observed to be closely associated with the shaft of the thick filaments in the 3D reconstructions from isolated thick filaments (*18, 19*). These interactions remain unexplored, but we predict that other myosin HCM mutations, not involved in the S1-S1, S1-S2, S1-MyBP-C and S2- MyBP-C interactions, may be involved in interactions between the folded complex and LMM.

While ~40% of HCM mutations are in human β-cardiac myosin, another ~40% are in MyBP-C, and a majority of those are truncation mutations, which may lead to loss of the protein and haploinsufficiency (*57, 58*). The considerations described here might suggest that at least the MyBP-C HCM truncation mutations are simply eliminating the protein, resulting in a loss of some of the interactions holding the sequestered complex together.

In summary, it is becoming increasingly clear that many myosin heads in the sarcomere are in a sequestered state held in equilibrium with actively cycling heads (Fig. 6). This equilibrium is shifted toward more active heads by phosphorylations of the myosin RLC and the M-domain of MyBP-C, providing fine-tuned control of cardiac function by adrenergic stimulation. Our hypothesis is that the equilibrium is also shifted toward more active heads by a majority of known myosin HCM mutations, and probably many MyBP-C HCM mutations as well, causing hyper-contractility of the heart. Modulating this equilibrium with small molecules may be the very best approach to new therapies for HCM, and dilated cardiomyopathy (DCM) as well, since the highly-tuned individual rate constants of the actin-activated myosin chemomechanical cycle could be left untouched, and only the number of functionally-accessible myosin heads (N_a_) for interaction with actin would be altered. Furthermore, small molecule therapeutics could be developed to interfere specifically with any one of the four types of intra- and inter-molecular interactions hypothesized here. Such an agent would also have the benefit of producing a limited maximum effect on heart function rather than completely inhibiting or over-activating the contractile apparatus.

**Figure 6.**
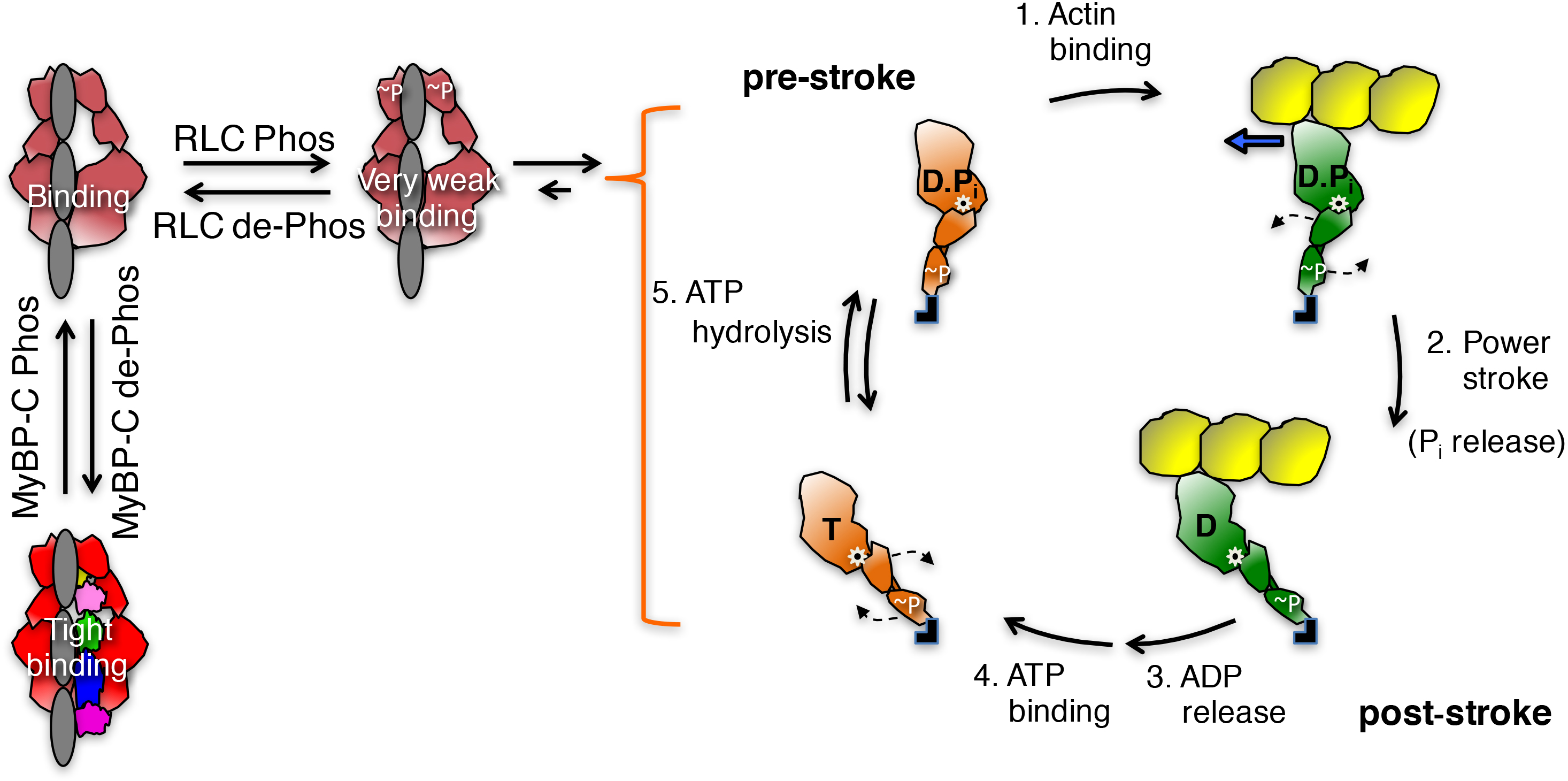
Schematic drawings of the actin-myosin chemomechanical cycle and hypothesized sequestered states of myosin heads. Steps: 1. The pre-stroke S1 (orange) with bound ADP (D) and phosphate (P_i_) binds to actin (yellow). 2. While bound to actin (green head), the lever arm swings to the right about a fulcrum point (black dot on white star) to the post-stroke position, moving the actin filament to the left (bold blue arrow) with respect to the myosin thick filament. 3. ADP release frees the active site for binding of ATP (T). 4. ATP binding weakens the interaction of the S1 to actin and frees the lever arm for cocking into the pre-stroke state. 5. ATP hydrolysis locks the head into the pre-stroke state. The heads in the cycle are phosphorylated (~P) on the RLC. Heads that are sequestered into a non-functional state (shades of red on the left side of the figure) are shown in three states: RLC phosphorylated and weakly bound to their S2 tail (grey), RLC de-phosphorylated and more tightly bound to their S2 tail, and complexed with de-phosphorylated MyBP-C and more firmly locked into the sequestered state.

## Acknowledgments

We thank Margaret Sunitha for help in homology modeling of both the human β-cardiac S1 and the full length MyBP-C. This work was funded by NIH grants GM33289 and HL117138 (to J.A.S.), the Shanta Wadhwani Centre for Cardiac and Neural Research, a Stanford Child Health Research Institute and the NIH-NCATS-CTSA grant #UL1 TR001085 (to D.V.T.). J.A.S. is a founder of Cytokinetics and MyoKardia and a member of their scientific advisory boards.

